# Safety and immunogenicity of an adjuvanted human onchocerciasis vaccine candidate, *Ov*MANE1: preclinical evaluation in mice model

**DOI:** 10.1101/2025.04.28.651093

**Authors:** Derrick Neba Nebangwa, Mary Teke Efeti, Sonia M. E. Momnougui, Cabirou Mounchili Shintouo, Robert Adamu Shey, Fidele Ntie-Kang, Rene Bilingwe Ayiseh, Stephen Mbigha Ghogomu

## Abstract

Onchocerciasis, caused by the filarial worm Onchocerca volvulus, remains a major public health challenge due to the limitations of ivermectin-based control strategies, thereby, highlighting the need for more innovative tools like vaccines. This study investigated the safety and immunogenicity of a novel multi-epitope chimeric antigen, OvMANE1 formulated with Freund’s adjuvant, in BALB/c mice. Following mice immunization at three time points of 2-week intervals, adjuvanted-OvMANE1 exhibited a promising safety profile, revealing neither any physical signs of toxicity nor behavioural abnormalities. Immunological assays showed significant increases in total IgG levels after the first (p = 0.0260) and final booster doses (p = 0.026). Interestingly, total IgG (p = 0.0086) and IgG1 (p = 0.0465) levels also increased significantly over the study period highlighting the ability of OvMANE1 to sustain humoral immunity. Moreover, cellular responses were significantly enhanced, with elevated leukocyte count (p = 0.0190) and increased lymphocyte activity (p = 0.0397) observed in the adjuvanted-OvMANE1 group compared to the control. Indeed, total leukocytes increased progressively from day 0 to day 39, with significant differences recorded in the test group between doses: day 0 vs. day 14 (p = 0.0043) and day 14 vs. day 28 (p = 0.0079). The pronounced production of relevant antibodies and induction of cellular immunity strongly suggests that the antigen can elicit mixed Th1/Th2 responses and antibody-dependent cellular cytotoxicity (ADCC) targeting O. volvulus L3 and/or other larval stages of the parasite. These results clearly show the emergence of OvMANE1 as a promising vaccine candidate against human onchocerciasis. However, further studies to evaluate the antigens protective potential in other animal species are required.

## Introduction

Human onchocerciasis, commonly known as river blindness, is one of the most devastating yet neglected tropical diseases (NTDs), caused by the filarial nematode *Onchocerca volvulus* through exposure to repeated bites of an infected *Simulium* blackfly that breeds around fast flowing rivers [1,2]. Onchocerciasis primarily manifests clinically through severe itching, disfiguring skin conditions, and visual impairments including permanent blindness, which has led to its recognition as the second leading infectious cause of global blindness, after trachoma [2]. Currently, *O. volvulus* is endemic in Africa, Central America, and South America with >99% of all cases occurring in sub-Saharan Africa [1]. According to the 2017 Global Burden of Disease Study, 14.6 million infected individuals were reported to have skin disease, while 1.15 million suffered from vision loss [3]. Recent WHO 2024 statistics, reports at least 249.5 million people in 28 countries requiring interventions to eliminate onchocerciasis with 18.8 million people needing preventive chemotherapy (PC) [2]. Due to its substantial impact on health, onchocerciasis has been a long-standing priority for international disease control groups like the Onchocerciasis Elimination Program for the Americas (OEPA) and the African Programme for Onchocerciasis Control (APOC). However, control efforts have primarily relied on two strategies: combating the *Simulium* fly vector through aerosol spraying and administering ivermectin as large-scale chemotherapy. While both methods have been used individually or in combination, these control strategies face several limitations [4]. These include insecticide and ivermectin resistance, high costs and logistical challenges, lack of ivermectin macrofilaricidal activity, ineffectiveness of sprays in certain areas, and the re-infestation of forests by black flies, hindering the complete elimination of onchocerciasis [5,6]. Moreover, the restriction on ivermectin usage in areas where onchocerciasis and loiasis co-exist, poses one of the most significant bottlenecks to onchocerciasis control.

Earlier in 2015, an international consortium launched a new global initiative, named The Onchocerciasis Vaccine for Africa (TOVA) with the key goal to advance the development of vaccine candidates meeting the desired target product profiles (TPP) for an onchocerciasis vaccine [7]. The TPP targets a preventive vaccine for children under the age of 5 who are restricted from taking ivermectin, or a therapeutic vaccine for both adults and children infected with onchocerciasis. Lustigman *et al*., (2018) reported TOVA’s process that led to the selection of several protein sub-unit vaccine candidates for further clinical development based on proven efficacy both *in vitro* and in animal models [8–12]. However, these antigens offer only partial protection [9,13]. Moreover, issues such as immune evasion, especially when addressing pathogens with immunomodulatory properties like *O. volvulus* [14], frequently lead to unsatisfactory results in human proof-of-concept clinical trials. Thus, there remains a dire need for more research addressing the design and development of more efficacious vaccines. Multi-epitope vaccines containing both B and T cell epitopes, covering various antigens expressed across multiple larval stages of parasitic worms have been suggested as innovative strategies for combating filarial infections [15]. These vaccines offer significant advantages, including the ability to induce a balanced Th1 and Th2-specific protective immune response, as well as risk minimization of adverse effects associated with unfavorable epitopes present in a full antigen sequence or whole parasite-based vaccines [16,17]. Novel multi-epitope vaccine candidates have demonstrated promising efficacy against other nematode infections, including lymphatic filariasis and trichinellosis [18,19].

Previously, we utilized immunoinformatics to design a multiple epitope vaccine candidate, *Ov*-MANE1 (**Fig 1A**), which incorporates relevant B and T cell epitopes derived from eight immunogenic peptides that were previously evaluated in preclinical studies [20]. In the process of cloning the chimeric antigen into a pMAL-c5X vector, an MBP tag was incorporated at the N-terminal of *Ov*MANE1 antigen. Addition of the MBP tag has several benefits including the facts that: i) MBP is known to enhance protein solubility and stability, improving the expression and purification of resulting fusion antigens, thus, facilitating large-scale pharmaceutical production [21]; ii) MBP additionally serves as a carrier protein by facilitating the targeting of antigens to antigen-presenting cells (APCs) that express mannose receptors, thereby enhancing the immunogenicity and ability of MBP-based antigens to stimulate Th1 responses and cytotoxic T lymphocytes [22]. This targeted antigen delivery improves the activation of both cellular and humoral immune responses, as evidenced by studies on mannosylated fusion proteins [22,23]. iii) Finally, mannose-binding protein is generally considered safe as it is a natural component of the innate immune system [23]. Overall, *Ov*MANE1 demonstrated remarkable antigenicity, supported by WormBase gene expression data, which confirmed that all peptides incorporated within its full *Ov*-MANE1 domain are expressed across every stage of the *O. volvulus* parasite [20]. This underscores its significant potential as both a prophylactic and therapeutic vaccine candidate, offering comprehensive protective coverage throughout the various stages of the *O. volvulus* life cycle [20]. Consequently, this study aimed at assessing the safety and immune response profile of *Ov*MANE1+Freund’s adjuvant in BALB/c mice model for further characterization of the antigen as a potential vaccine candidate against onchocerciasis. The findings demonstrated pronounced production of relevant IgG antibodies and cellular components, including lymphocytes. This strongly suggests that the antigen has the capacity to elicit mixed Th1/Th2 responses and antibody-dependent cellular cytotoxicity (ADCC) targeting *O. volvulus* L3 and/or other larval stages of the parasite.

**Fig 1:**
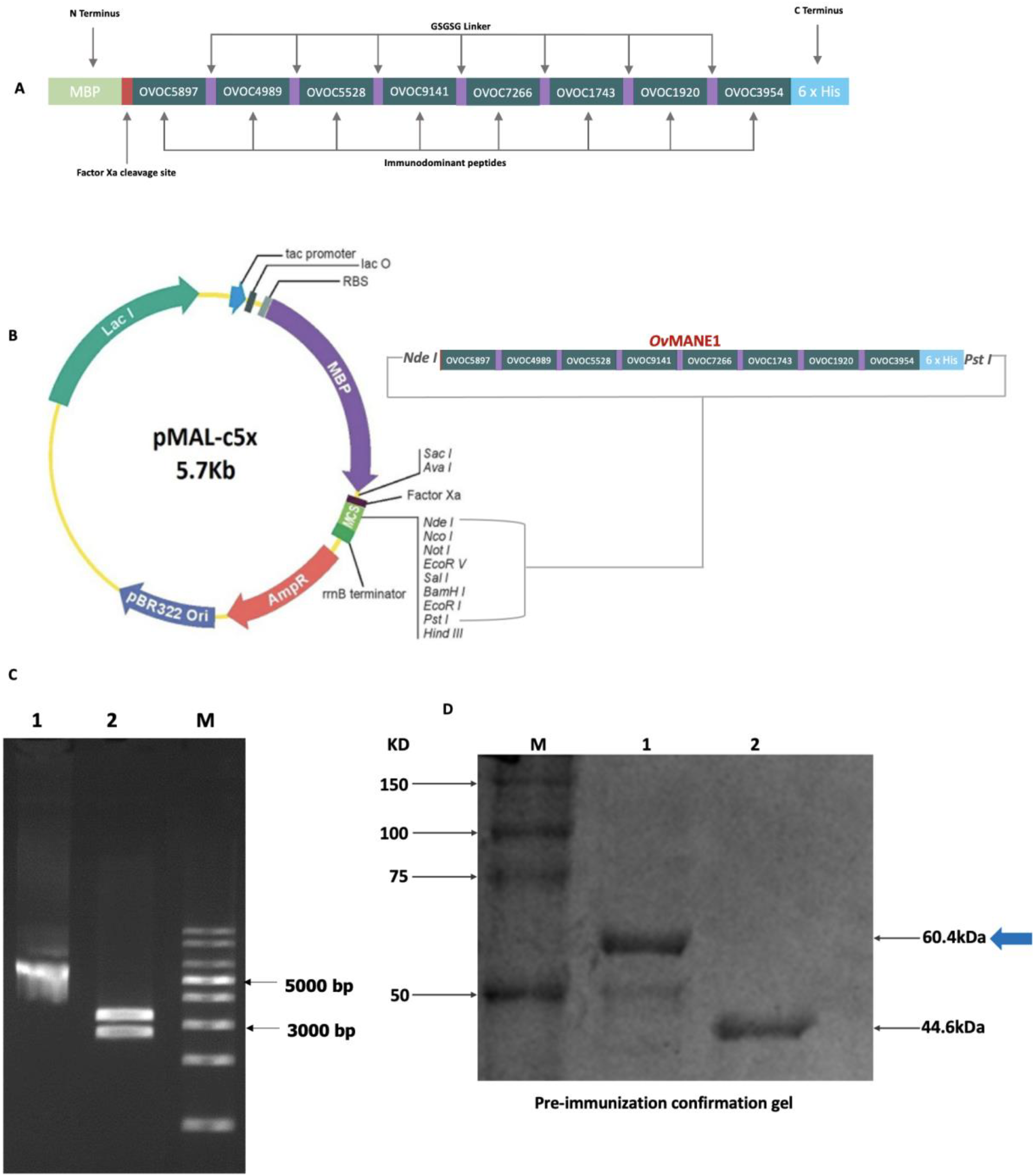
Schematic representation, expression, purification, and detection of *Ov*MANE1. A) Representation of *Ov*MANE1 (560 amino acid residues) consisting of eight immunodominant peptides (IDPs colored navy green) fused with GSGSG linkers (purple stripes) to form the *Ov*MANE1 domain; linked at the N-terminal to a mannose-binding protein (MBP) (MBP colored light green) and at the C-terminal to a 6x histidine tag (colored light blue) aiding with downstream purification and characterization [20]. B) Expression plasmid map indicating OvMANE1 gene inserted in frame with the MBP-tag at the N-terminal between NdeI and Pst restriction sites. C) Digestion gel of the recombinant pMAL-c5x plasmid coding for ovMANE1. Expected fragment sizes include for lane 1 = OvMANE1 single digest with MluI – 6096 bp and for lane 2 = OvMANE1 double digest with MluI and HindIII – 3348 bp and 2748 bp. D) SDS-PAGE confirmation of the integrity of purified *Ov*MANE1 antigen prior to mice immunization. Lane 1= *Ov*MANE1 with band at approximately 60.4 kDa, and Lane 2=MBP band at approximately 44.6 kDa. M: ladder (molecular weights). Blue arrow indicates the positions of *Ov*MANE1. Proteins were resolved with 12.5% Sodium Dodecyl Sulfate–Polyacrylamide Gel Electrophoresis (SDS-PAGE) gel stained with Coomassie blue.

## Results

### Expression, purification, and confirmation of *Ov*MANE1 chimeric antigen

*Ov*MANE1 chimeric antigen previously designed and reported by us as a promising biomarker for developing a diagnostic test for human onchocerciasis [20] is made up of eight *O. volvulus* immunodominant peptides derived from the following antigens: OVOC5897, OVOC4989, OVOC5528, OVOC9141, OVOC7266, OVOC1743, OVOC1920 and OVOC3954. The rationally screened immunodominant peptides were fused using flexible GSGSG linkers to enhance the accessibility of the epitopes to antibodies and improve protein folding (**Fig 1A**). The chimeric antigen was cloned into pMAL-c5X vector **(Fig 1B)**, and then successfully expressed, and purified as previously reported in our previous study [20]. WormBase gene expression data indicated that all eight peptides making up the *Ov*MANE1 domain were expressed in all the parasite stages of *O. volvulus*. The addition of an MBP tag to *Ov*MANE1 upon expression in a pMAL-c5X vector serves to enhance *Ov*MANE1’s solubility, stability, and large-scale pharmaceutical production [21] while improving antigen targeting to antigen-presenting cells (APCs) for robust Th1 and cytotoxic T-cell responses. Before mice immunization, the purified antigen was assessed on a 12% polyacrylamide gel to verify its integrity (**Fig 1D**). As expected, a distinct *Ov*MANE1 band was identified at approximately 60.4 kDa, confirming its integrity for use in downstream mice immunization experiments.

### Qualitative safety assessment of adjuvanted-*Ov*MANE1 antigen

Assessment of adverse effects following immunization of mice with the adjuvanted-*Ov*MANE1 antigen primarily involved qualitative monitoring of physical signs of toxicity and behavioral changes after immunization. Aside from mild erythema (redness of the skin), no other injection site reactions, such as swelling or skin rashes, were noteworthy in any of the groups. Behavioral patterns were also monitored post-immunization, with no events such as unusual lethargy or hyperactivity detected. Additionally, eating and drinking behaviors, including appetite and hydration levels, were normal across all groups, and no abnormal social interactions, like aggression or withdrawal patterns observed. These results suggest that *Ov*MANE1 formulated with Freund’s adjuvant might be well-tolerated in BALB/c mice.

### ELISA analysis demonstrates substantial IgG antibody responses to adjuvanted-*Ov*MANE1

Measuring total IgG levels reflects how effectively the humoral immune system is stimulated by antigens in this context. Previous studies have shown that IgG1 sub-type specifically plays a crucial role in protecting against parasitic infections [9,24]. Accordingly, an indirect ELISA was performed to assess total IgG and IgG1 responses to *Ov*MANE1, using sera from immunized mice across the three treatment groups. The findings indicated a significant rise in antibody titers on day 25 following the first boost (day 14; *p* = 0.0260) and on day 39 after the final boost (day 28; *p* = 0.026) in the adjuvanted-*Ov*MANE1 group compared to the adjuvant control group (**Fig 2A**). Additionally, within the group of mice receiving *Ov*MANE1 formulated with Freund’s adjuvant, total IgG levels steadily increased over time (days 0 to 39) with successive immunization doses, demonstrating a significant (*p* = 0.0086) difference in antibody levels between the baseline and post-final booster dose (day 39) (**Fig 2B**). For IgG1, while the adjuvanted-*Ov*MANE1 group had higher titres post-final booster dose compared to both the adjuvant and PBS control arms, the difference was not significant (**Fig 2C**). Interestingly, a detailed analysis of the *Ov*MANE1 formulated with Freund’s adjuvant group revealed a statistically significant increase (*p* = 0.0465) in IgG1 responses over time, demonstrating a consistent upward trend across successive immunizations, from the initial prime dose to the final booster dose (**Fig 2D**). These findings show clear development of relevant IgG antibodies, suggesting the capacity of the antigen to elicit antibody-dependent cellular cytotoxicity (ADCC) targeting *O. volvulus* L3 and/or other larval stages of the parasite, a key protective mechanism against onchocerciasis.

**Fig 2:**
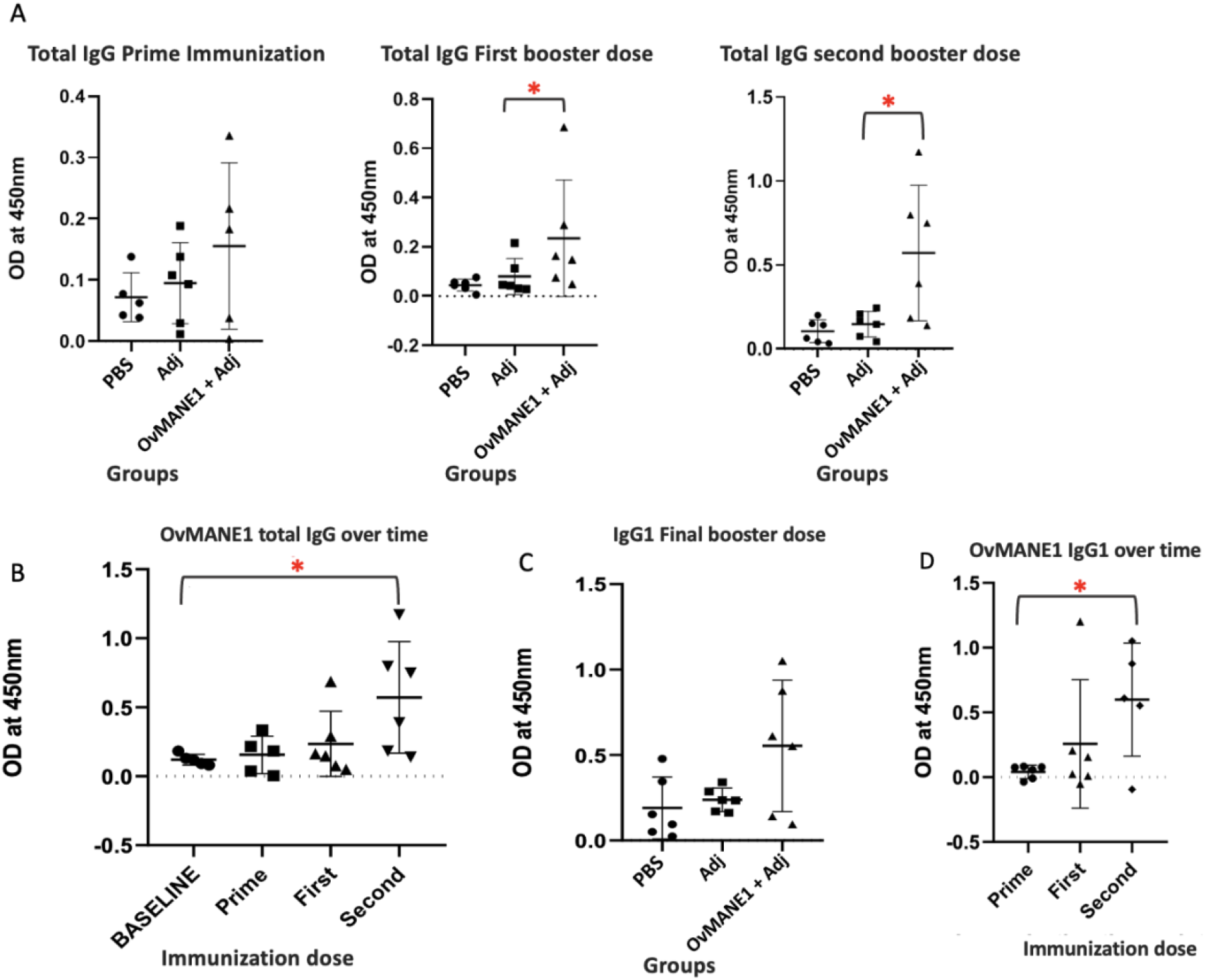
IgG antibody responses to *Ov*MANE1+Freund’s adjuvant relative to control groups and across time. The purified antigen was used to coat microtiter plates. Microtiter plates were blocked and incubated with mice sera from the three treatment groups, obtained at days 11, 25, and 39 after the prime, first boost, and second booster immunization doses respectively, at a dilution of 1:27,000 followed by incubation with goat anti-human IgG peroxidase conjugate at a dilution of 1:5000 as the secondary antibody. The plates were then revealed using TMB, the optical density (OD) was read at 450 nm and OD values were plotted for the different groups. A) Analysis between treatment groups showed higher levels of total IgG in the *Ov*MANE1+Adjuvant group at all points of immunization, compared to the controls (PBS and adjuvant-only groups). B) Analysis within the adjuvanted-*Ov*MANE1 group showed a significant (*p* = 0.0086) increase in total IgG response over the study period with successive immunization doses from baseline (day 0), prime immunization (day 11), first boost (day 25) and second boost (day 39). C) While IgG1 titers were higher in the test group than the controls, the difference was not significant after the final booster dose (day 39). However, D) Analysis within the treatment group showed a significant difference (*p* = 0.0465) in IgG1 levels between the prime and second booster doses. PBS=phosphate buffer saline, Adj=Adjuvant, Prime=prime immunization dose, First=first booster injection, Second=second booster dose.

### Leukocyte responses indicate that adjuvanted-*Ov*MANE1 functions as a potent immunogen

Leukocyte (white blood cells) counts serve as a standard marker for evaluating the cellular immune potential of antigens or adjuvant formulations in experimental studies [25]. Therefore, to evaluate the cellular responses to *Ov*MANE1+Freund’s adjuvant at different immunization time points, the total and differential white blood cells count was performed by Giemsa stain microscopy. Overall, a statistically significant rise in total leukocyte count was recorded following the second booster dose administered on day 28 (*p* = 0.0190) in the adjuvanted-*Ov*MANE1 group when compared to the PBS control group (**Fig 3A**). Moreover, in the group of mice immunized with *Ov*MANE1 formulated with Freund’s adjuvant, total white blood cell count showed a progressive increase over the study period (days 0 to 39) with each successive immunization dose, reflecting significant differences in leukocyte levels between the post-prime dose (day 0) and post-final booster dose (day 39) (*p* = 0.0043); as well as between the post-first booster dose (day 25) and post-final dose (day 39) (*p* = 0.0079), as illustrated in **Fig 3B**.

**Fig 3:**
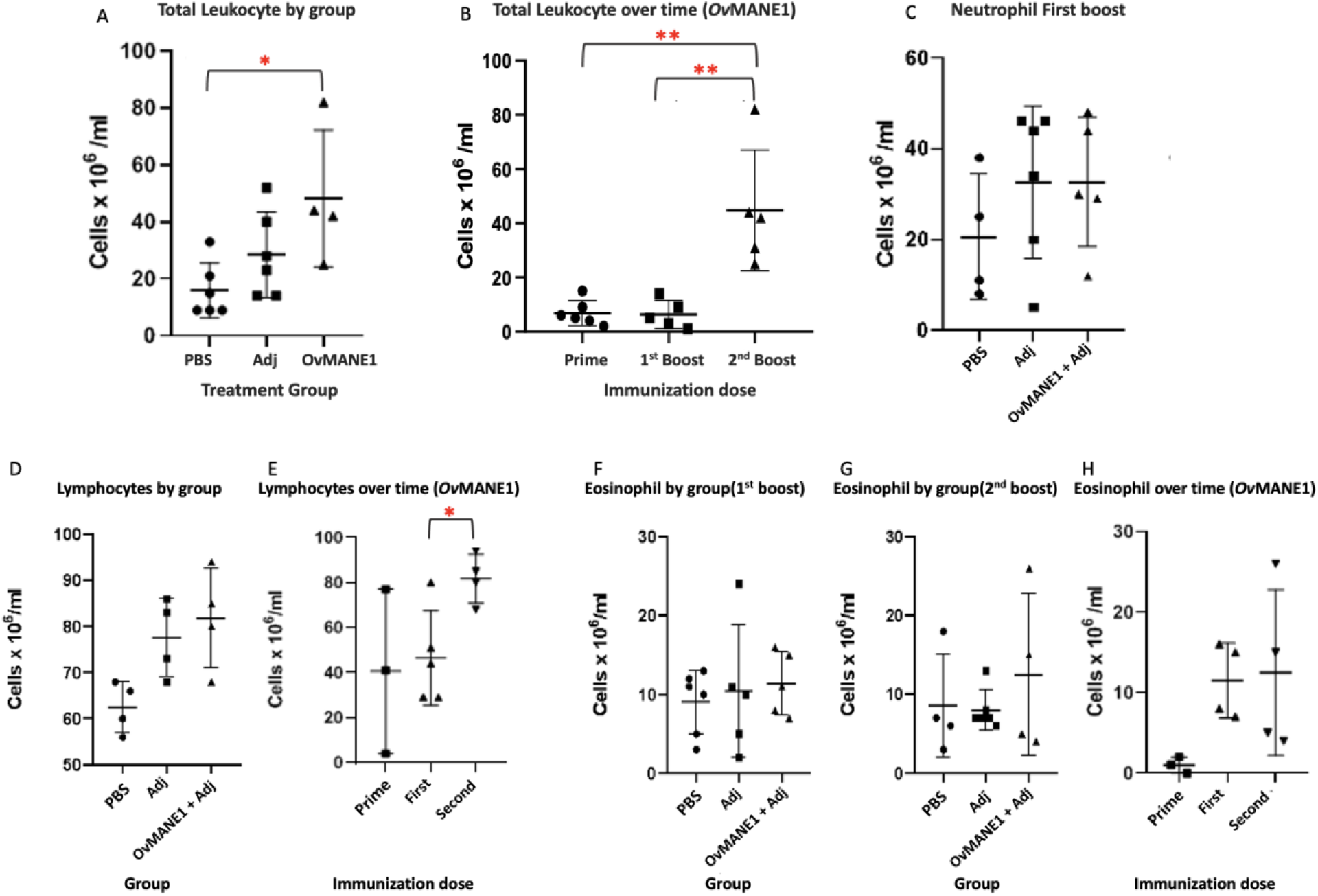
White blood cell responses to adjuvanted-*Ov*MANE1. Mice whole blood samples were collected into EDTA-coated tubes and diluted with Turk’s solution in a 1:20 ratios and a drop of the mixture loaded on a hemocytometer, mounted under a 40x objective lens of a microscope and from the large corner squares on the grid patterns, white blood cells count was recorded and compare between the groups at every immunization time point. A) Total leukocyte responses between treatment groups post-final booster immunization dose. A significant increase (*p* = 0.0190) was observed in the adjuvanted-*Ov*MANE1 group compared to the PBS control group, after the second boost (day 39). B) Total leukocyte responses within the *Ov*MANE1+adjuvant group across immunization time points. A general significant increase in total leukocyte levels was observed in the *Ov*MANE1 group, between the prime (day 0) and post-second boost (day 39, p = 0.0043) and between the post-first booster dose (day 25) and post-second booster dose (day 39, *p* = 0.0079). **Differential white blood cell assessment:** Mice EDTA-whole blood samples (5 µl) were used to prepare a thin blood smear on microscopic slides and stained with 10% Giemsa using standard Giemsa staining protocol. Differential white blood cell counts, focusing on neutrophils, lymphocyets, and eosinophils were performed manually by examining slides under 100x oil immersion light microscope, counting at least 100 white blood cells. C) Neutrophil levels post-first booster immunization timepoint for the different treatment groups. The neutrophil levels were higher in the adjuvanted-*Ov*MANE1 group compared to the PBS control, although not significant. D) Lymphocyte levels post-final booster immunization across the various treatment groups. E) Lymphocyte responses within the group of mice immunized with the *Ov*MANE1 formulated with Freund’s adjuvant across immunization time points. A significant increase was observed between the post-first booster dose and post-final boost (*p* = 0.0397). Eosinophil responses across the treatment groups F) post-first booster immunization dose and G) post-final immunization. In both cases, the adjuvanted-*Ov*MANE1 group exhibited higher eosinophil counts compared to the controls. H) Distribution trend in eosinophil levels for the *Ov*MANE1+adjuvant group across immunization timepoints with progressive increase from prime dose on day 0 to post-final immunization dose on day 39. PBS=phosphate buffer saline, Adj=Adjuvant, Prime=prime immunization dose, First=first booster injection, Second=second booster dose

To further determine the specific cellular responses to the adjuvanted-*Ov*MANE1, differential leukocyte levels, with a focus on neutrophils, lymphocytes, and eosinophils, were evaluated among the different treatment groups at different immunization time points. The result showed that the *Ov*MANE1+Adjuvant group had more neutrophils and lymphocytes induced than the PBS and adjuvant control groups post-final (day 39) immunization, although the difference in lymphocyte count between the test and control groups was not significant (**Fig 3C-D**). However, the data indicated a significant increase (*p* = 0.0397) in lymphocyte counts within the adjuvanted-*Ov*MANE1 treatment arm over time, between the post-first booster dose (day 25) and post-final booster dose (day 39) as shown in **Fig 3E**.

Furthermore, eosinophil responses were evaluated across the different treatment groups and across successive immunizations over time. The results indicate that the adjuvanted-*Ov*MANE1 group exhibited higher eosinophil counts following both the first (day 25) and second (day 39) booster doses compared to the control groups, although the differences were not statistically significant in either case (**Fig 3F-G**). Likewise, for the mice immunized with *Ov*MANE1 formulated with Fruend’s adjuvant, sharp and mild elevations in eosinophil levels were noted between the post-prime (day 11) and post-first (day 25) booster dosing periods, as well as between the post-first (day 25) and post-second (day 39) booster dosing intervals, respectively, over the study timeline. However, these increases were not significant (**Fig 3H**). Nonetheless, these findings collectively demonstrate that adjuvanted-*Ov*MANE1 elicits a strong and sustained leukocyte-mediated immune response, underscoring its potential as a promising immunogen.

## Discussion

River blindness continues to pose a major challenge in developing countries like Cameroon, where community-directed treatment with ivermectin, the main control tool against the infection, faces notable limitations such as ivermectin resistance and the lack of macrofilaricidal drugs [26,27]. As a result, there is an urgent need for alternative intervention strategies, including prophylactic vaccines to combat the disease, supporting the WHO’s 2030 global onchocerciasis elimination agenda [28]. Our research, thus, aimed to evaluate the capability of a novel multi-epitope chimeric antigen, *Ov*MANE1, to safely elicit suitable humoral and cellular immune responses following immunization of BALB/c mice at three different time points of two-week intervals. The assessment of adjuvanted-*Ov*MANE1 immunization in mice revealed no significant physical signs of toxicity nor behavioural abnormalities, suggesting its promising safety profile and suitability for clinical development. On the other hand, analysis of the humoral responses demonstrated a significant increase in total IgG and IgG1 antibody levels in the adjuvanted-*Ov*MANE1-immunized group compared to controls (**Fig 2**). Similarly, whole, and differential leukocyte analyses reveal that *Ov*MANE1+adjuvant induced a strong and sustained cellular immune response in mice, underscoring its potential as an effective immunogen (**Fig 3**).

*Ov*MANE1 recombinant chimeric antigen was successfully designed and produced from synthesis of the conceptual gene construct through expression to purification, as previously described in our previous study [20]. Similar approaches using multi-epitope chimeric antigens have demonstrated strong immunogenic potential in other studies, such as in phase I clinical trials targeting *Plasmodium falciparum* antigens [29,30], thus, recommending the promising profiles of such multi-peptide vaccine candidates. The design of *Ov*MANE1 incorporated an MBP motif at its N-terminal to improve the antigen’s solubility and stability, with the goal of simplifying and enabling effective large-scale pharmaceutical manufacturing [21] while facilitating the targeted delivery of *Ov*MANE1 segment to antigen-presenting cells (APCs), thereby driving strong Th1 responses and cytotoxic T-cell activation [22,23]. Adjuvanted-*Ov*MANE1 demonstrated exceptional antigenicity in enzyme-linked immunosorbent assays. A key strength of the antigen is relies on the fact that it consists of eight immunodominant epitopes expressed across all the life cycle stages of *O. volvulus* parasite [20], emphasising its potential as both a prophylactic and therapeutic vaccine against river blindness.

Indeed, mice immunization with *Ov*MANE formulated with Freund’s adjuvant was well-tolerated, as we observed no physical signs of toxicity and behavioural changes including lethargy, hyperactivity, and abnormal social interactions. Nonetheless, the mild erythema observed at mice injection sites across all treatment groups is consistent with immune activation responses typically associated with tissue injury due to injection site skin damage, as well as antigen-adjuvant combinations, as reported by Kool *et al*., (2008) [31]. However, the stable appetite and hydration levels witnessed across all mice groups throughout the study duration further strengthen the case for *Ov*MANE1 as a safe antigen for clinical development.

*Ov*MANE1 also induced potent antibody responses, as evidenced by a significant rise in total IgG antibody titres following the first (*p* = 0.0260) and second (*p* = 0.0260) booster doses (**Fig 2A-B**), compared to adjuvant control. Similarly, the gradual increase in antibody levels over time, equally, underpins the effectiveness of Freund’s adjuvanted-*Ov*MANE1 in stimulating sustained humoral immune activation. These findings align with previous research illustrating that multi-epitope antigens can elicit strong and durable humoral immunity in murine models [13,30]. The significance of total IgG and IgG1 in protecting against onchocerciasis has been highlighted [32] and supports the relevance of the elevated IgG and IgG1 antibody responses to *Ov*MANE1 detected herein. Therefore, the significant and steady increase in total IgG (*p* = 0.0086) and IgG1 (*p* = 0.0465) level observed between the baseline and final booster immunizations further supports capacity of the adjuvanted-*Ov*MANE1 to prime and enhance humoral immunity, making it a promising candidate for onchocerciasis vaccine development. These results correspond with numerous studies highlighting the cytophilic effect of IgG antibodies against *O. volvulus* parasites in both murine and non-human primate immunization challenge experiments [9,11].

Our findings also indicate that *Ov*MANE1+adjuvant effectively induces cellular immune responses, as the total leukocyte counts following booster immunizations significantly increased (*p* = 0.0190) than the control (**Fig 3A**). This observation corresponds with prior research demonstrating similar patterns of leukocyte proliferation in response to protective filarial antigens [33,34]. The progressive rise in total white blood cell counts throughout the immunization schedule, with significant differences between first-boost and post-final booster (*p* = 0.0043) (**Fig 3B**), suggests successful immune priming and memory response induction, and so aligns with similar findings from other recombinant helminth vaccines [35]. Although neutrophil and lymphocyte proliferation in mice immunized with adjuvanted-*Ov*MANE1 was not significantly elevated compared to the controls, notable increases were observed relative to the PBS control (**Fig 3C-D**). Moreover, lymphocyte responses within the *Ov*MANE1+adjuvant group demonstrated significant temporal elevations (*p* = 0.0397) (**Fig 3E**), and eosinophil counts exhibited marked increases following booster doses (**Fig 3F-H**). However, the absence of statistically significant differences in eosinophil levels compared to the control groups warrants further investigation. These trends are consistent with observations from promising vaccine candidates for related filarial parasites [34,35], where early cellular responses preceded stronger adaptive immunity. The sustained leukocyte-mediated immune activation elicited within the adjuvanted-*Ov*MANE1-immunised mice group might present *Ov*MANE1 as a potential vaccine candidate against human onchocerciasis, though additional research on long-term immunity and protective efficacy is required to confirm its full prospects.

Certain mechanisms underlying acquired protective immunity against *O. volvulus* infection in humans (**Fig 4**) have been examined, and so revealed that protection is linked to the ability to elicit mixed Th1/Th2 responses targeting *O. volvulus* L3 and/or other larval stages of the parasite [36,37]. Furthermore, that effective anti-helminth immunity through antibody-dependent cell-mediated cytotoxicity (ADCC) against L3 is facilitated by cytophilic IgG antibodies including IgG1 and IgG3; as well as cytokine release [10,36]. In this research, we recorded marked yet consistent increases in total IgG levels and significant IgG1 levels (*p* = 0.0465) (**Fig 2**), along with substantial proliferation of overall cellular responses, including a significant rise in lymphocytes over time and notable rise in neutrophil and eosinophil counts when comparing the adjuvanted-*Ov*MANE1-vaccinated mice group to the PBS control (**Fig 3**).

**Fig 4:**
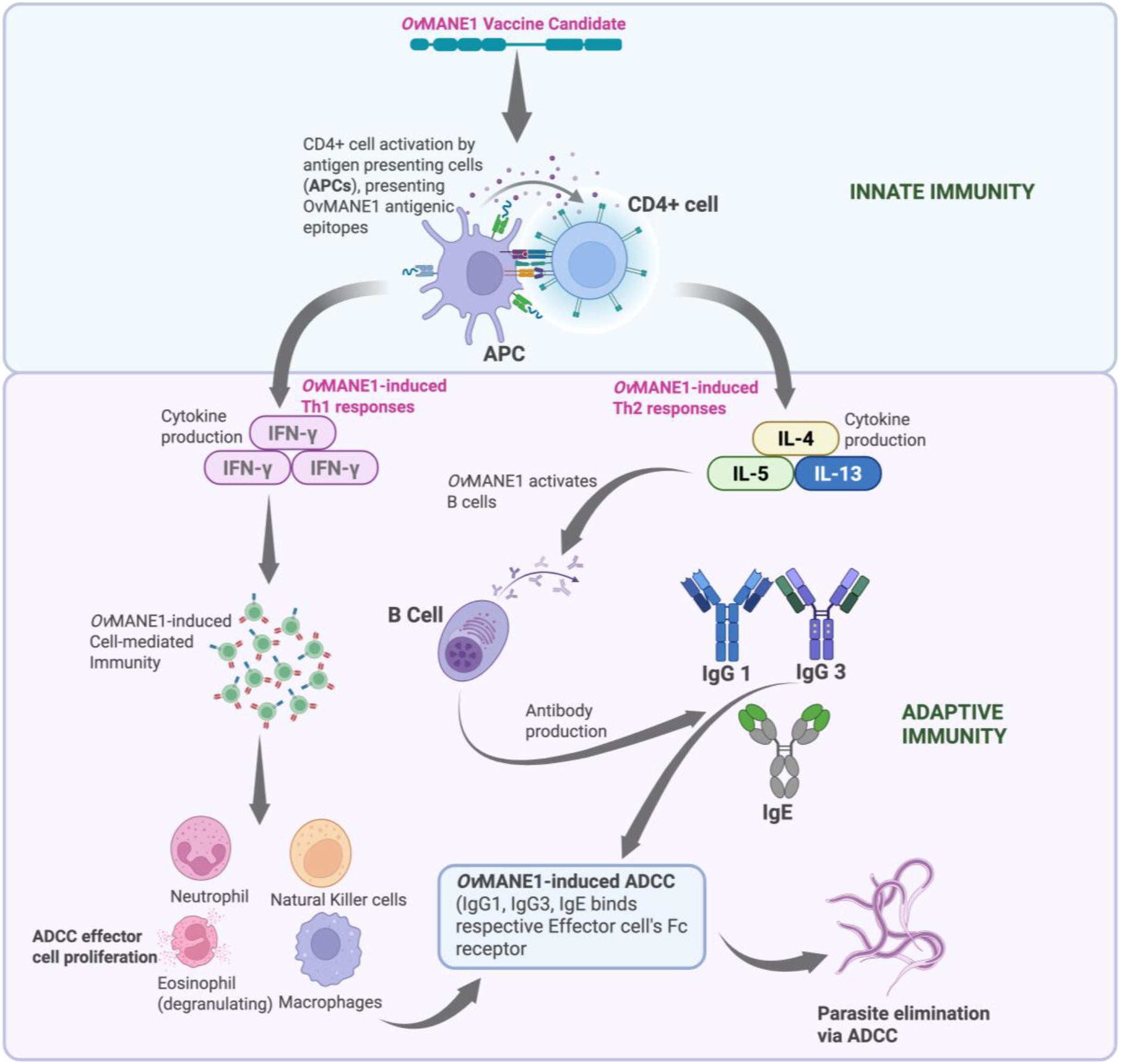
Schematics of *Ov*MANE1’s potential protective immune mechanisms. The figure illustrates the immunological mechanisms that might be activated by the *Ov*MANE1 vaccine candidate, based on the immune response profiles observed in this study, where marked levels of immune cells (neutrophils, lymphocytes, and eosinophils) and IgG antibodies (total IgG + IgG1) are observed. It emphasizes the potential innate and adaptive immune responses to the antigen. The upper section highlights the activation of antigen-presenting cells (APCs), which process and present *Ov*MANE1 epitopes to CD4+ T cells (lymphocyte), inducing distinct Th1 and Th2 responses. *Ov*MANE1-induced Th1 responses would potentially stimulate cytokine production, particularly interferon-gamma (IFN-γ), which supports cell-mediated immunity and activates effector cells (neutrophils and eosinophils with marked proliferation observed herein), potentially contributing to antibody-dependent cellular cytotoxicity (ADCC) through expression of their respective families of Fc receptors for antibody binding. In contrast, potential *Ov*MANE1-induced Th2 responses would drive the production of interleukins such as IL-4, IL-5, and IL-13, promoting humoral immunity and B-cell activation. Through adaptive immunity, B cells then produce immunoglobulins (total IgG and IgG1 with marked increases in titers as observed in this study) in response to Th2 cytokine stimulation. These antibodies would then bind to effector cells via Fc receptors, completing the ADCC pathway and enhancing parasite elimination.

**Fig 5:**
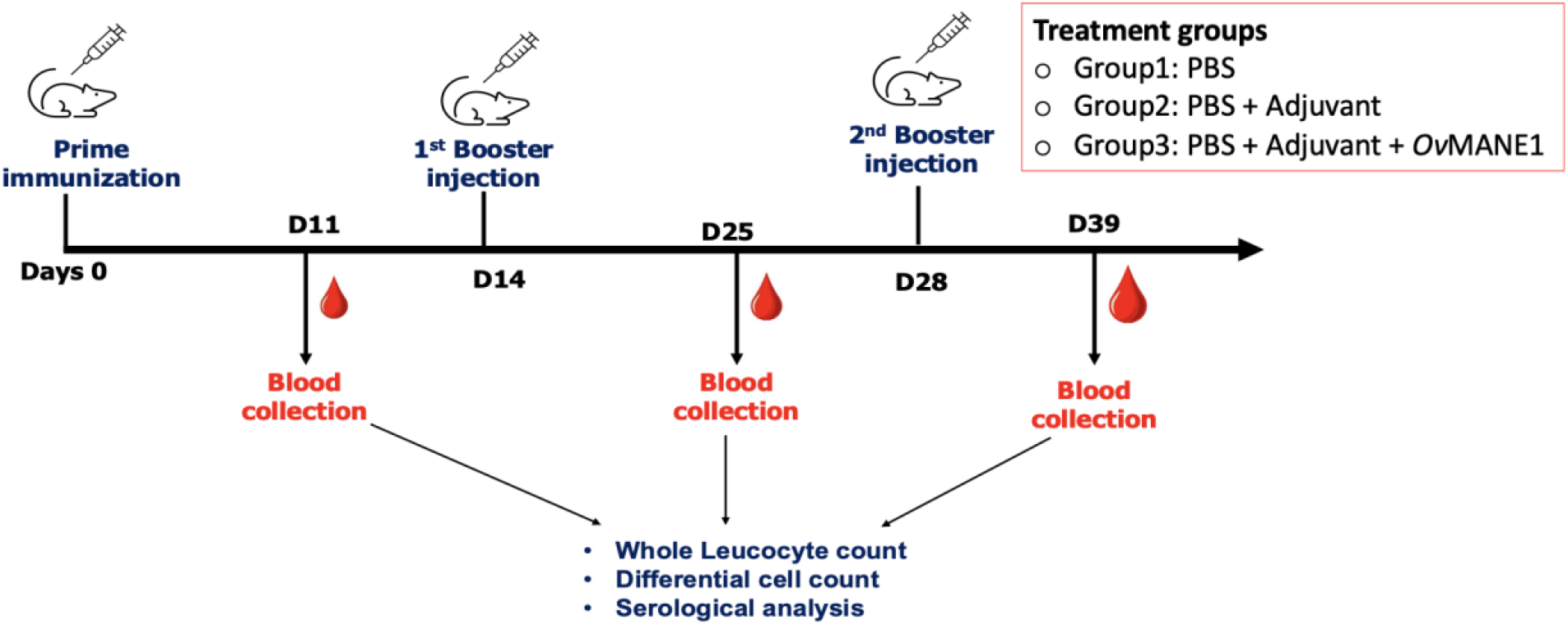
Mice immunization and sample collection plan. BALB/c mice were subcutaneously immunized thrice at two-week intervals. Whole blood was collected by retro-orbital bleeding at various time points for whole leucocyte counts, differential cell counts and serological analysis. D11-39 represents days 11 to 39. PBS: vehicle negative control group. Adjuvants were Freund’s complete adjuvant for prime dose and Freund’s incomplete adjuvant for booster doses.

Consequently, the evident formation of IgG antibodies and cellular elements, including neutrophils and eosinophils following immunization, strongly indicates the ability of *Ov*MANE1 to elicit antibody-dependent cellular cytotoxicity (ADCC) (**Fig 4**). Indeed, earlier studies suggest that neutrophils and eosinophils can facilitate ADCC through Fc receptor expression, thereby amplifying their cytotoxic efficacy [38,39], while also accentuating their involvement in ADCC, particularly neutrophils’ capacity to bind IgG antibodies via its Fc receptor. Additionally, the significant proliferation of T lymphocytes (**Fig 3E**) contributes to anti-helminth immunity through cytokine production (particularly Th2 responses including IL-4, IL-5, and IL-13) which supports eosinophil function and antibody production. Eosinophils can then target helminths through ADCC-related IgE-mediated mechanisms, releasing their toxic granule proteins [40]. Increased total IgG1 levels are indicative of a Th2-skewed response and associated cytokines like IL-4 and IL-5, aligning with findings by Du *et al*., (2020) [41], while lymphocyte proliferation upon immunization suggests overall immune activation, indicative of proliferation of Th1 cells which serve as key drivers of cellular immunity, and associated with cytokines like IFN-γ [42].

Despite these encouraging data, our study is not without limitations. For instance, the research was restricted to murine models, which might not fully represent the intricate immune responses seen in humans. Furthermore, a key methodological limitation arose from the reliance on light microscopy to evaluate cellular immunity to the antigen. This approach introduced a degree of subjectivity in visual interpretation, which could have resulted in confirmation bias. The method was also unable to distinguish between CD4+ and CD8+ cells. Moreover, consistent blood sampling of mice using standard retro-orbital bleeding protocol proved challenging. Consequently, concerns regarding potential harm to the animals and limitations associated with capillary tube blood clothing rendered it impractical to collect samples from some mice at the predefined time points. This limitation resulted in missing data points for antibody and cellular responses for some mice across treatment groups. Nevertheless, future research employing advanced techniques (e.g blood sampling and flow cytometry) are required to achieve a more precise and detailed characterization of immune responses to *Ov*MANE1.

In summary, our study provides strong evidence underpinning the potential of the novel multi-epitope chimeric antigen, *Ov*MANE1, as a promising vaccine candidate against onchocerciasis. Immunization of BALB/c mice elicited significant increases in both humoral and cellular immune responses, including evident increases in total IgG and IgG1 antibody levels, as well as notable leukocyte proliferation, particularly involving lymphocytes, neutrophils, and eosinophils. These findings suggest that *Ov*MANE1 could effectively protect against *Onchocerca volvulus*, hence contributing to the World Health Organization’s 2030 goal of onchocerciasis elimination. Future investigations could include detailed studies in other small animal species and non-human primates to assess the safety, immunogenicity, and efficacy of *Ov*MANE1 in much more detail and context. Additionally, data from animal challenge experiments with *O. volvulus* or sister species such as *O. ochengi* parasites are needed to ascertain the protective immune mechanisms of *Ov*MANE1. Understanding the durability of the induced immunity and elucidating the mechanisms driving the immune response to this antigen will be crucial for its clinical development and use.

## MATERIAL AND METHOD

### Ethical consideration for animal studies

Ethical clearance for the use of mice models was obtained from the university of Buea Institutional Animal Care and Use Committee (IACUC) with reference number (UB-IACUC Nº 02/2021). The validated ethical consideration protocols were adhered to throughout the study to ensure the humane treatment of the animals.

### Animal model and grouping

This study involved BALB/c mice purchased from the Jackson Laboratory (Bar Harbor Maine, USA), aged 6–8 weeks, weighing approximately 16–27g. Mice were housed in micro-isolator boxes in standard laboratory conditions (temperature: 23 ± 2°C and 12-hour light/dark cycle) and provided with *ad libitum* access to food and water. The mice were randomly assigned to three treatment groups: (1) control group receiving phosphate-buffered saline (PBS), (2) adjuvant control group receiving Adjuvant + PBS, and (3) test group receiving *Ov*MANE1 antigen + Adjuvant + PBS (**Table 1**). Each group consisted of 6 mice. All protocols involving mice were approved by the Institutional Animal Care and Use Committee of the University of Buea.

**Table 1:**
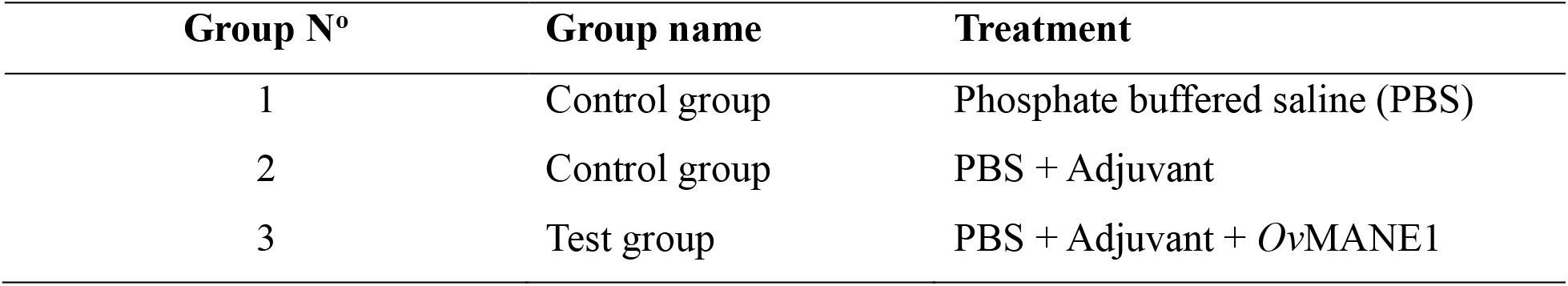
Mice immunization treatment groups.

### Collection of mice baseline data

The individual weights of the mice were measured using an electronic weighing scale and mice were tagged by serial numbering from one to eighteen. Blood samples were collected through retro-orbital bleeding following standard protocol [43] into 0.5mL EDTA coated microcentrifuge tubes for preliminary whole leucocyte and differential cell count. Blood samples were aliquoted into dry 0.5mL Eppendorf tubes, centrifuged at 13000 rpm for 15 min, and serum collected into sterile 0.5mL Eppendorf tubes and stored at -20 ^o^ C for downstream serological analysis.

### Mice immunization

The total antigen and vehicle volumes for immunization were prepared per group based on the description on table 2. The solutions were prepared for three injection doses; priming, first boost and second booster injections following standard protocol [43]. Freund’s complete adjuvant (CFA) was used for the priming, while Freund’s incomplete adjuvant (IFA) was used for the two booster injection doses with antigen and vehicle volumes as described in Table 2. To prepare the adjuvanted antigen solution, CFA or IFA adjuvants were added to *Ov*MANE1 antigen solution (cloned, expressed, and purified as previously reported by us [20]) and the mixture vortexed. After an eight-day mice acclimatization period, a 25gauge syringe was used to deliver a 150 µL injection containing respective treatment doses to each mouse. Injections were administered subcutaneously at the nape. The initial prime dose was administered on day 0, followed by two booster doses on days 14 and 28 (**Fig 4**). One hundred microliters of blood was collected by retro-orbital bleeding using standard protocols [43] at baseline and on days 11, 25, and 39 post-immunization for immunological assays. Additionally, serum samples were obtained at similar timepoints for serological analysis (Fig 1). Qualitative assessment of adjuvanted-*Ov*MANE1’s safety was performed post-immunization. Mice were observed qualitatively for physical and behavioral signs of toxicity and adverse side effects *ad libitum* over the duration of the study. Behavioral monitoring encompassed observations of activity levels (checking for lethargy or hyperactivity), feeding and drinking patterns, as well as social interactions among the mice including potential aggressive behavior or withdrawal patterns.

**Table 2:**
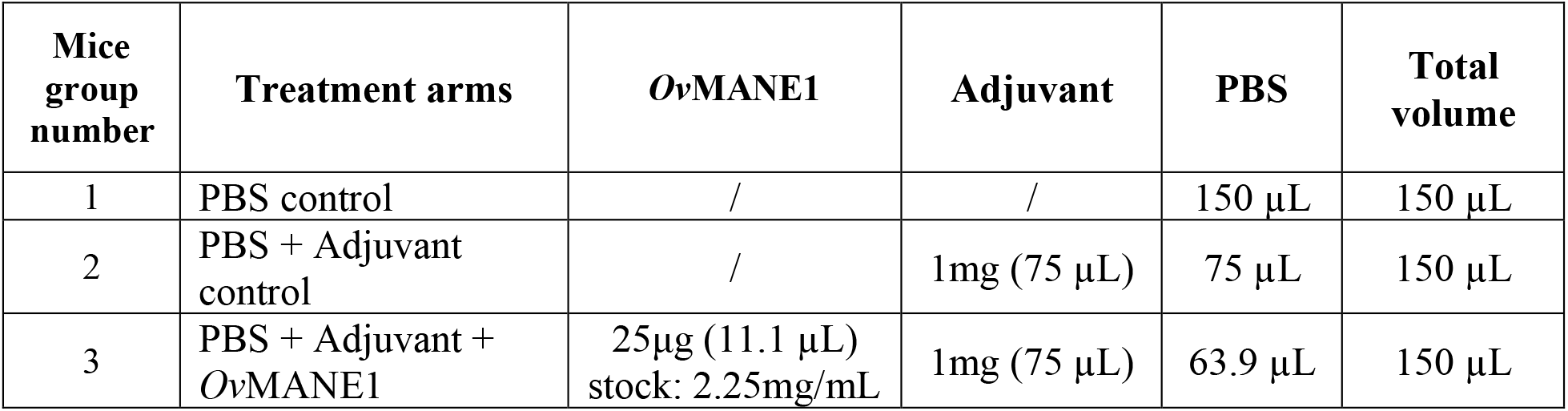
Distribution of volume of immunization doses per mice group.

### Whole leukocyte and differential white blood cell count

The whole leukocyte count was performed to provide an overview of the general immune cell response to *Ov*MANE1. Turk’s fluid was prepared locally with Gentian violet, acetic acid and distilled water according to the standard preparation protocol [44] and blood samples previously collected in EDTA-coated tubes were diluted into Turk’s solution in a 1:20 ratio. A drop of the mixture was then loaded on a hemocytometer and mounted under a 40x objective light microscope lens and observed according to standard leukocyte enumeration protocol. Leukocytes were counted and recorded using the formula:

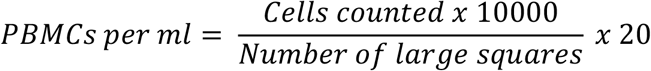

To understand the specific components of the immune responses, differential white blood cell count was also performed. About 5 µl of mice EDTA-whole blood samples were used to prepare a thin blood smear on microscopic slides and stained with 10% Giemsa using standard Giemsa staining protocol (S. S. College, Jehanabad’s protocol of Enumeration of WBC – Total Leukocyte Count (TLC)). Differential white blood cell count was performed manually by examining slides under a 100x oil immersion light microscope, counting at least 100 white blood cells. Lymphocyte neutrophils, and eosinophil cell types were identified and differentially classified.

### Antibody detection by enzyme-linked immunosorbent assay

The ELISA protocol used by Arumugam *et al*. (2016) with slight modification was used to detect the presence of antibodies (IgG and IgG1) induced by *Ov*MANE1 in the serum of immunized mice. Briefly, MaxiSorp^TM^ 96-well microtiter plates (Nunc, Roskilde, Denmark) were coated with 50 µl/well of 2 µg/ml of *Ov*MANE1 in 0.5M carbonate buffer (pH 9.4) at 4°C overnight. Plates were then washed thrice with 0.1% PBS-Tween-20 (PBS-T) after a 5 min interval and blocked with 200µl/well of 3% non-fat milk for 2 hours at room temperature. The plates were then washed thrice with PBS-T, and individual serum samples diluted at 1:27,000 in 1% non-fat milk and 50 µl serum samples added to corresponding wells. Plates were incubated at room temperature for 2 hours and washed thrice with PBS-T. Goat anti-human peroxidase conjugated IgG (Merck Millipore, Billeria, MA) was prepared in a 1% non-fat milk solution at a 1:5000 dilution, and 50 µl was added into each well. Plates were incubated for an hour before being washed thrice again with PBS-T. Finally, 50 µl of the TMB-KPL substrate was added to each well and the colour was allowed to develop for 10 minutes at room temperature. The reaction was stopped by adding 50 µl of 3M HCl per well. The optical densities were recorded using a microplate reader at 450 nm and antibody levels estimated. All washing and antibody dilutions were done in the wash buffer (0.1% PBS-Tween-20).

### Statistical analysis

All data were recorded and processed using Microsoft Excel 2013. Statistical analyses were performed with GraphPad Prism 8.0 software (GraphPad Software, La Jolla, CA, USA). Antibody titers were compared using the Mann Whitney t-test (u-test) for nonparametric data analysis. ANOVA (Analysis of Variance) was used to identify significant differences between groups, while a *p*-value less than 0.05 was considered as the threshold of statistical significance for all parameters.

## Supplementary Materials

All data included in paper

## Author Contributions

**Conceptualization –** DNN, RBA, RAS, and SMG

**Data curation –** DNN, MTE, and SMEM

**Formal analysis –** DNN, MTE, SMEM, CMS, and RBA

**Funding acquisition –** DNN and SMG

**Investigation –** DNN, MTE, SMEM, CMS, RAS, and RBA

**Methodology –** DNN, CMS, RAS, and RBA

**Project administration –** SMG, RBA, and FNK

**Resources –** RAS, RBA, FNK, and SMG

**Software –** FNK, DNN, and RBA

**Supervision –** RBA, FNK, and SMG

**Validation –** RBA, FNK, and SMG

**Visualization –** DNN, MTE, SMEM, CMS, and RBA

**Writing – original draft –** DNN and MTE

**Writing – review & editing –** All authors

## Institutional Review Board Statement

Use of mice and the experimental procedures performed in this study were reviewed and approved by the IACUC committee of the University of Buea, Cameroon.

## Acknowledgments

This work was supported by a grant from the International Society for Infectious Diseases and the European Society of Clinical Microbiology and Infectious Diseases awarded to DNN and SMG. We express our gratitude to Professor Mark R. Sanderson of Imperial College London for supporting DNN through tuition and research fee supplements. Additionally, we acknowledge Mrs. Eyong Etaka Judith, Mr. Arrey Etta Augustine, Mrs. Gemuh Blendin Sirri, and Mr. Feko Thadde Fredy for their valuable research assistance during the mice studies. The funders had no role in the study design, data collection and interpretation, or the decision to submit the work for publication.

